# An Interpretable Deep Learning Approach for Biomarker Detection in LC-MS Proteomics Data

**DOI:** 10.1101/2021.02.19.431935

**Authors:** Sahar Iravani, Tim O.F. Conrad

## Abstract

Analyzing mass spectrometry-based proteomics data with deep learning (DL) approaches poses several challenges due to the high dimensionality, low sample size, and high level of noise. Additionally, DL-based workflows are often hindered to be integrated into medical settings due to the lack of interpretable explanation. We present DLearnMS, a DL biomarker detection framework, to address these challenges on proteomics instances of liquid chromatography-mass spectrometry (LC-MS) - a well-established tool for quantifying complex protein mixtures. Our DLearnMS framework learns the clinical state of LC-MS data instances using convolutional neural networks. Based on the trained neural networks, we show how biomarkers can be identified using layer-wise relevance propagation. This enables detecting discriminating regions of the data and the design of more robust networks. One of the main advantages over other established methods is that no explicit preprocessing step is needed in our DLearnMS framework.

Our evaluation shows that DLearnMS outperforms conventional LC-MS biomarker detection approaches in identifying fewer false positive peaks while maintaining a comparable amount of true positives peaks.

**Code availability:** The code is available from the following GIT repository: https://github.com/SaharIravani/DlearnMS

## 1 Introduction

LIQUID chromatography-mass spectrometry (LC-MS) based proteomics allows analysis and quantification of complex protein mixtures. This technique quantifies the components based on their physio-chemical properties and their molecular mass which yields a LC-MS map with two orthogonal dimensions chromatographic retention time (RT) and mass to charge ratio *m*/*z*. Due to the precise and fast analysis, the LC-MS technique has been widely used in high-throughput proteomics applications, such as biomarker detection, disease diagnosis/prognosis, or drug target identification [1], [2], [3]. The main difficulties of analysing LC-MS maps, however, lies in their properties: they are high-dimensional, typically highly complex, and contain a high level of noise. This makes it for example very challenging to detect biomarkers from raw LC-MS maps of proteins [1], [4]. The idea of biomarker detection - sometimes also called feature selection - is to identify proteins by which a specific medical condition can be determined. Thus, biomarkers are differentially abundant single peaks specified by *m*/*z* and RT on a raw LC-MS map.

### 1.1 Related work

Many of the well established tools for LC-MS biomarker discovery – such as MsInspect [5], MZmine 2 [6] or Progenesis [7] – are organized in multiple (often three) main stages. They usually begin with a pre-processing stage that commonly includes noise reduction, RT alignment [8], [9], [10], data normalization [11], data filtering [12], baseline correction, and peak grouping. This is followed by a second stage that involves peak detection. Here, informative areas within the LC-MS maps are extracted. This is done in multiple ways – MsInspect, for example, identifies peaks using wavelet decomposition, MZmine 2 applies a deconvolution algorithm on each chromatogram to detect peaks, and Progenesis uses a wavelet-based approach, but this time in such a way that all relevant quantitation and positional information are retained. Other well known frameworks include: XCMS [13] where the peak detection step is addressed by a pattern matching approach on overlaid extracted ion chromatograms with Gaussian kernels; AB3D [14] which iteratively takes the highest intensity peak candidates and heuristically keeps or removes neighboring peaks to form peptide features; MSight [15] which adapts an image-based peak detection on the generated images from LC-MS maps; and MaxQuant [16] in which a correlation analysis involving a fit to a Gaussian peak shape is applied. A common design in all these methods is that they all dependent on some kind of pre-defined model for signal detection. One of the main benefits of the Deep Learning approaches is that the model is not pre-defined, but rather learned from the input data through the training phase. We will show later that this is indeed beneficial if it is combined with an interpretation phase.

After this second stage an analysis of the detected peaks is done to identify the actual biomarkers. However, two main problems that arise often during the aforementioned steps are (1) that low-intensity peaks are lost and (2) many parameters need to be tuned for these methods to perform well, e.g. by adjusting them for new data sources.

In this paper, we present a novel approach for biomarker detection in LC-MS maps that does not need explicit preprocessing steps. Our deep learning based approach implicitly learns the needed transformations and can identify biomarkers with an overall improved performance compared to the mentioned traditional approaches.

### 1.2 Deep Learning for Proteomics Analysis

The success of deep learning (DL)-based methods, which have been replacing state-of-the-art methods in many fields [17], [18], [19], have also entered the field of proteomics data analysis already some time ago. Notable examples are: DeepIso [20], which combines a convolutional neural network (CNN) with a recurrent neural network (RNN) to detect peptide features; DeepNovo [21] and DeepNovo-DIA [22] that use DL-based approach (CNN coupled with RNN) for peptide sequencing on data-dependent acquisition (tandem mass spectra) and data-independent acquisition MS data, respectively; pDeep [23] adapts the bidirectional long short term memory for the spectrum prediction of peptides; and DeepRT [24] employs a capsule network to predict RT by learning features of embedded amino acids in peptides.

Despite the current successful approaches, most of these studies are empirically driven, and are lacking a justifiable interpretation foundation [25]. Moreover, as machine learning (ML) and DL have been rapidly growing also for real-world applications, a concern has emerged that the high precision accuracy may not be enough in practice [26]. Rather, interpretation and understanding of the made decisions is important for robustness, reliability, and enhancement of a system. On top of it all, supervised data-driven biomarker detection models require annotated data at the peak level which is in most cases rather expensive or even infeasible to acquire. To address these challenges, in this paper we leverage deep learning interpretability to understand and analyze LC-MS proteomics data which requires just the instances class labels for training, rather than expensive peak annotations.

### 1.3 Interpretation of Deep Neural Networks

Methods for interpreting Deep Neural Networks (DNNs) provide information about what makes a network arrive at a certain decision. These methods can roughly be divided into four categories: (1) the *function* analysis that explains DL model itself through gradient and shows how much changes in input pixels affect the output [27], [28], (2) the *attribution* method that interprets the output of the model and explain which features and to what extent contribute to the model’s output [29], [30], [31], (3) the *signal* method that tries to find patterns in inputs on which the decision is based on [32], [33], [34], and (4) the *perturbation* analysis that calculates the importance of features through measuring the effect of perturbing the elements of inputs on the output [35], [36], [37]. The application of DNN explanation employing perturbation analysis has previously studied in metabolomics [38]. However, permutation analysis is not computationally feasible for high-throughput LC-MS analysis. Among three other interpretation categories the out-performances of *attribution* analysis has been demonstrated in [25] on MALDI-TOF MS data. In this study, therefore, one of the methods in *attribution* category called layer-wise relevance propagation (LRP) [31] is employed for interpretation of the model predictions of LC-MS proteomics. To guarantee the trustworthiness of the LRP explanation in our feature selection task we analyse the sensitivity of the interpretations in terms of their repeatability, reproducibility, and their robustness.

**Fig. 1.**
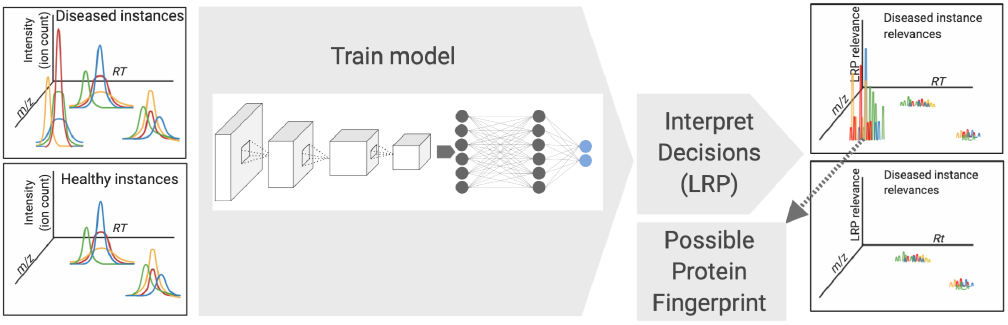
Overview of our DLearnMS approach foTr discovery of disease related biomarkers.

### 1.4 DLearnMS: a novel approach for biomarker detection

In this paper we present DLearnMS, a biomarker detection approach based on interpretable deep learning to allow analyzing and – ultimately – understanding LC-MS data. The basic idea is as follows: Given two groups of LC-MS samples (say, healthy and diseased), a convolutional neural network (CNN) is trained, and the learned configuration is interpreted through the layer-wise relevance propagation (LRP) technique. We use the result from the interpretation step to identify the areas in the input-data that play a crucial role for differentiating the two groups. This is analysed further to firstly verify the robustness of the network and improve the network architecture and secondly detect the differentially abundant peaks as biomarkers. Our biomarker detection model benefits from optimizing on class labels rather than expensive annotations at peak levels. Since high-quality labeled datasets are not widely available, we suggest a method to tune the network architecture using synthetically generated data through performing systematic series of experiments and quantitatively measuring the interpretations. We evaluate the proposed model also on real-world data and demonstrate the superiority of DLearnMS compared to conventional biomarker detection frameworks. One of the major advantages here is that our method does not depend on the otherwise necessary preprocessing steps. Nevertheless, LC-MS preprocessing approaches e.g., [39], [40], could be potentially added to DLearnMS framework for further improvement. The stability of the detected biomarkers including the reproducibility and robustness are examined through cross-validation strategies. Finally, we discuss the shortcomings of conventional ML models for analysing raw LC-MS data classification and interpretations. Our contributions in this paper lies in the combination of the following triad:

A. We present an interpretable Deep Learning (DL)-based approach that can identify biomarker candidates from high-throughput LC-MS proteomics data. One of the main advantages is that the method does not need (potentially) expensive peak level annotations.
B. We show how to use layer-wise relevance propagation as an interpretation technique for deep learning networks and how this can be used for feature identification in this context.
C. We demonstrate how to tackle the scarcity and sparsity of labeled LC-MS data by using synthetically generated data and evaluate the improvements compared to only using experimental data.

## 2 Designing the Model

Let 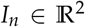 for *n* = 1, …, *N* be a set of LC-MS maps with *O_n_* ∈ {0, 1} as the assigned class labels (e.g. the respective medical conditions). Each (*x*, *y*) pair on *I* where *x* = *m*/*z* and *y* = *RT*, contains the ion-counts of the LC-MS map. The aim of biomarker detection is to find the smallest subset of 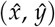 pairs where the ion-counts are differentially abundant between conditions 0 and 1. Our strategy is to design a CNN architecture, modeled as a function *f*, to classify LC-MS samples into two classes, and learn from the prediction behavior to detect 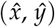 pairs. Mathematically speaking, a CNN with *L* layers can be abstracted as *f*(*I*) = *f_L_* ○ … ○ *f*_1_(*I*) where each layer is a linear function followed by an elementwise non-linear activation, such as the rectified linear unit function (ReLu [41]). The power of CNN prediction comes from combining many layers, which at the same time makes it complex and consequently difficult to interpret. The layerwise relevance propagation (LRP) method [31] uses the layered structure of the neural network to interpret the predictions. The network is assumed to be fully trained in order to use LRP, and the predictions are redistributed backward layer-by-layer to give a score to all the input features. A feature 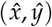 will be attributed with *strong relevance*, if the function *f* is sensitive to the presence of that feature. The relevance values of all (*x, y*) pairs form the matrix of relevances 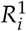 is known as a heatmap. The goal is to adapt this information for verifying the network predictions of medical conditions and learn form the network behaviour to find the most relevant attributions associated with this these condition.

### 2.1 Classification Model and Interpretation

The first step is to design a robust classification CNN for the LC-MS samples of two classes where we are interested in the differences. A CNN is usually characterized by the depth and width of the layers. Depth refers to the number of layers, and width determines the number of filters. We train multiple types of networks with different width and depth based on standard structures like variants of ResNet [42] and also tailored (or customized) structures. We observe that training very deep networks like ResNet32 on the LC-MS data (both synthetic and real data) leads to overfitting. The better performance of our shallower network compared to the very deep networks can intuitively be explained by the local dependent characterization of the peaks on the LC-MS map. Very deep networks capture both the local - gained by reach feature representation - and global dependencies - gained by large receptive fields. Therefore, very deep networks may learn some global patterns irrelevant to the data information but relevant to the noise, such as quantification calibration error in the data acquisition. Apart from the depth of the network, we observe that changing a few layers on the architecture of the customized network has not change the training and testing accuracy and loss. To decide keeping or removing these layers, one may select a network with fewer learnable parameters to decrease the computational cost. Whereas, one my select a network with more learnable parameters to increase the capacity and a better generalization accuracy. Our strategy to select a proper network architecture is however to leverage CNN interpretation. We quantitatively compare the interpretations of the network predictions with different architectures, and select the one whose predictions are aligned the most with the actual differences between the two groups. To obtain the interpretations we employ LRP method using Eq. (1). Applying LRP on the network’s prediction of given input *I_n_* highlights the important parts of *I_n_* through redistributing the neuron score backwards through the layers until the input layer and assigns a relevance to each element of the input. Eq. (1) shows a rule for redistributing the relevances known as LRP.*ϵ*.

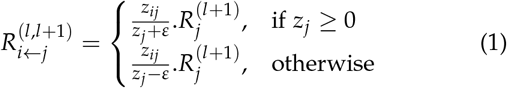

where 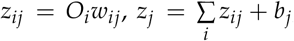, and *O_j_* = *g*(*z_j_*). *g* is a non-linear activation function, and *w_ij_* defines the weight that connect the neuron *j* in layer *l* to the neuron *i* in layer *l* + 1. Other redistribution rules to control the flow of positive and negative relevances include LRP.*αβ* and LRP.*z* [31]. All rules at each step must hold such that 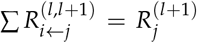, which means all relevance values that flow into a neuron at layer *l* + 1 flow out towards the neurons of the layer l. All Relevances, 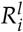, are calculated for *l* = 1, …, num_layers progressively from last layer, layer after layer, until the input layer is reached and yield 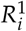. Please see [31] for more details. 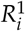 for *i* = (*x*, *y*) demonstrates how much pixel (*x*, *y*) - representing *m*/*z* and RT - contributes to the decision making. We choose a network whose 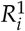 highlight the differences between the classes the most. As quantitatively assessing the interpretations requires the annotations at the peak levels, this experiment is performed on large synthetically generate data. The detail explanation on selecting the network architecture is delayed to Section 3.

### 2.2 Feature Selection

Once the network architecture has been selected, we employed the network to learn representation of real data and to discover the discriminating peaks from its interpretation. Our assumptions to use the interpretation for biomarker discovery is the reproducibility and robustness of the interpretations, which are justified later in Section 3.5. Considering offsets, the presence of noise, and different peak indices on the samples, we are interested in interpreting the decisions on statistics of the whole training-set. We take the mean of LC-MS samples belonged to the diseased class *D* and healthy class *H*, separately. Each mean is given to the trained network *f* and the predictions are interpreted by LRP function. This results in two matrices of diseased relevance values 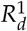 and healthy relevance values 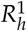.

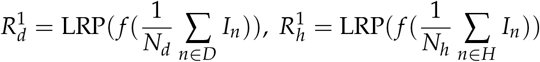

where *N_d_* and *N_h_* are the number of samples in diseased and healthy classes, respectively. The spatial location of peaks, however, on LC-MS map are widely distributed, where we estimate the exact location of peaks by finding their index with maximum intensity within a predefined window. To this end, first select the peak with strongest relevances on 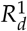. Then, the neighbor’s relevances in the window are set to zero. We iterate this process until all the high-intensity relevances are covered. The selected peaks are distinguished as biomarkers if corresponding indices on 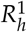 are attributed non-negative relevances. We will discuss the effect of incorporating 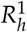 along with 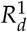 in Section 3.3.

To extract the biomarkers of a test sample, the sample is fed to the trained network to be classified. The peaks are selected locally from LRP interpretation similar to selecting the peaks from training samples. These peaks are distinguished as biomarkers if corresponding indices on 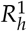 are existed and attributed non-negative on 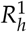.

## 3 Model Parameter Tuning

As our DLearnMS feature selection is built on top of a trained deep convolutional classifier, our aim in this section is to select a network architecture that is more reliable and robust to be the basis of our feature selection model. To this end, we interpret different trained architectures as heatmaps and assess which heatmap is aligned more with the discriminating regions of the data. We will show that although the variation in some layers results in very small differences in accuracy, their interpretation focus towards discriminating peaks differ. To assess the interpretation, however, we need the annotations at the peak level. Since the annotation at the peak level in large amount is too expensive or infeasible to acquire, this experiment is performed on a synthetically generated dataset. In the following, we first introduce how the synthetic LC-MS dataset is generated, and then describe how the network architecture is tuned through assessing their interpretation quantitatively.

### 3.1 LC-MS Data Simulation

LC-MS consists of two levels of separations. First, a protein solute (mobile phase) passes through a chromatography column (stationary phase), which effectively separates the components based on the chemical affinity and weight. RT measures the time taken from the injection of the solvent to the detection of the components. Second, each component is ionized and scanned through a mass spectrometer that generates a mass spectrum (MS). Each MS scan measures *m*/*z* values of charged particles and peak intensities. Stacking all MS scans on top of each other forms a three-dimensional data whose *x*, *y*, and *z* axes are *m*/*z* values, RT, and ion-count intensities, respectively.

To generate the synthetic LC-MS dataset, two groups of samples representing healthy and diseased classes are simulated using UniPort human proteome dataset [43]. The healthy class contains 20 peptides. Two peptides that are independent from the peptides in the healthy samples are added to the peptides in healthy group to form the diseased group. As a results, there are 20 and 22 peptides in healthy and diseased group. The two extra peptides in diseased group define the biomarkers (discriminating features) that we intend to detect on LC-MS map. Investigating such differences is the basis of diagnosis of different biological conditions and disease treatment, e.g., measuring the concentration level of cardiac troponin that enters in the blood soon after a heart attack, or measuring thyroglobulin, a protein made by cells in the thyroid, which is used as a tumor marker test to help guide thyroid cancer treatment.

We use OpenMs [44] and TOPPAS [45] to generate LC-MS samples and convert them into images. The width, height, and pixel intensities of images present *m*/*z*, RT, and ion-count intensity, respectively. It should be noted that the images still represent the raw data. The only difference between the matrix of raw data and the converted images is that the ion-count intensity range in raw data is scaled to [0,255]. The dataset contains 4000 samples of each group. 10% of each group is left out for testing, and the rest is used for training and validation.

### 3.2 Interpretation Assessment Metrics

Lets now introduce the metrics we selected to evaluate the capability of interpretation heatmap 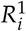 on reflecting the discriminating regions. The metrics should be representative of the percentage of true-positive (TP) and false-positive (FP) peaks. Therefore, we consider intersection over union (IOU), precision, and recall metrics defined as follows:

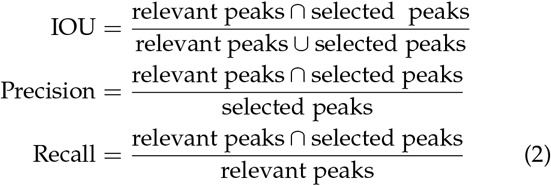

where the relevant peaks and selected peaks are ground-truth and predicted peptides peaks. To extract the ground truth on synthetic data, the mean of the images in the diseased group is subtracted from the mean of the images in the healthy group and the absolute value of the resulting is taken. The result contains all discriminating peaks and is referred to as ground-truth image (GTI). This is identical to average of several replicas of the spike-in peptides. We apply a threshold, *γ_gt_*, on the GTI to ignore small perturbation generated by LC-MS quantification error. As previously described in Section 2.2 since the spatial location of peaks is distributed widely, we restrict our attention to the peaks with the highest intensities in local region. To this end, first the index of the highest intensity value on GTI is selected. Then, the surrounding peaks in the window of w and h are set to zero. Next, this process is iterated until all the high-intensity peaks are covered. We refer to the resulting as ground truth peak map (GTPM). The selected peaks in Eq. (2) are extracted similar to GTPM from the LRP relevances and form prediction peak map (PPM). The metrics of Eq. (2) can be rewritten as follows:

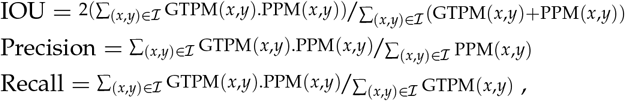

where 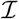 covers the entire range of (*m*/*z*,RT) values.

### 3.3 Network Architecture Selection

Up to this point, we explained the specification of the synthetic data, and introduced the metrics for interpretation assessment. We will now discuss how we choose and improve the network architectures including the number of FCL, CL, and MPL through interpretation assessment on synthetic data. Our experiment on the synthetic data shows that changing these parameters in a variation presented in Table 1 does not change the classification accuracy while their interpretation move significantly towards the discriminating peaks for making decisions. To show this effect, these networks are separately trained, and their interpretations are assessed using IOU, Precision, and Recall in Table 1.

**TABLE 1.** Network architecture selection through interpretation assessment. This table shows the effect of adding fully connected layers (FCLs), convolutional layers (CLs), max-pooling layers (MPLs) on focusing of the network on the discriminating peaks for decision making. The parameters are tuned according to the intersection over union (IOU), precision, and recall. The effect of incorporating the interpretation of diseased samples’ mean (*R_d_*) and the interpretation of healthy samples’ mean (*R_h_*) on peak detection is also demonstrated.

| # CL | # MPL | #FCL | Samples | IOU | Precision | Recall |
| --- | --- | --- | --- | --- | --- | --- |
| 6 | 4 | 2 | $R_d$ | 0.3975 | 0.3814 | 0.4149 |
| 6 | 4 | 1 | $R_d$ | 0.5006 | 0.4513 | 0.5621 |
| 6 | 4 | 1 | $R_d, R_h$ | 0.6177 | 0.6188 | 0.616 |
| 4 | 3 | 1 | $R_d$ | 0.6599 | 0.5985 | 0.7353 |
| 4 | 3 | 1 | $R_d, R_h$ | 0.7008 | 0.6756 | 0.7281 |
| 4 | 1 | 1 | $R_d$ | 0.7165 | 0.6171 | 0.8441 |
| 4 | 1 | 1 | $R_d, R_h$ | **0.8501** | **0.8554** | **0.8448** |

We see in our experiments that networks with higher values of interpretation assessment metrics - IOU, precision, and recall-are more generalized due to the fact that these networks know on which part of the data look for the reason of distinguishing a sample in one class from others and less biased toward irrelevant regions; therefore, it is more likely to act as the same on an unseen data. According to the research in DL field, exploiting deeper networks are recommended for better generalization as they offer richer representation. Contrary to our results (see Table 1), the deeper networks (more CL and FCL layers) show less reliance on the discriminating peaks. As a result, among the networks with the same accuracy performance, the one with four CL, one FCL, and one MPL reach the best interpretation performance. Hence, with more confidence we can say that the network distinguishes the samples according to the regions of the data that are truly discriminating. This is also the way how we as human would make classification decisions. Therefore, we can hope for more generalization performance for the real world instances where sometime strong perturbations are possible and clearly hard to predict. (The classification performances of designed network on simulated LC-MS data and real LC-MS data are depicted in supplementary material).

### 3.4 Interpretation Importance Across Different Classes

After selecting the network architecture, we now explain the effect of incorporating the interpretation of the healthy predictions along with the interpretation of the diseased predictions on reducing the FP peaks. In Section 3.2, we described in detail how prediction peak map (PPM) is calculated through LRP relevance values. As a recap, to estimate relevance values on the training set, we calculate the mean of the diseased samples, run the trained network on this mean, and calculate the relevances. By convention, positive relevance values are the evidence of existing relevant peaks that are belong to the respected class. Therefore, in our study, positive relevance values on the interpretation of diseased class have been associated with the discriminating peaks. We now aim to experimentally show that with the information from interpretation of healthy instances we can reduce the FP peaks that are highlighted with the interpretation of the diseased instances. This is because the positive relevances in the interpretation of the healthy instances can be explained as the absence of diseased relevant peaks, or presence of healthy relevant peaks. In our study, since all the discriminating peaks are appeared in diseased class, the positive relevances of healthy group is just explained as the absence of diseased relevant peaks. Accordingly, in our feature selection pipeline, the indices of high-ranked relevances in the diseased group are selected as biomarkers only if the corresponding indices in the healthy group attribute non-negative relevances. The results of this study are shown in Table 1, in which the interpretation column is assigned with *R_d_*, *R_h_*. As it is apparent, IOU and Precision that are both directly affected by FP in their denominator, have considerably improved.

As a result of architecture selection and parameter tuning, the feature selection performance has been improved from 40% to 85% shown in Table 1. Hence, our verified DL network architecture has four CL, one MPL after the second CL, and one FCL on top of the network as the prediction layer. We use the interpretation of this network for biomarker detection as it has been described in Section 2.2.

### 3.5 Interpretation Sensitivity analysis using Cross-Validation

The sensitivity analysis of deep learning interpretation methods has recently gained attentions with the aim of addressing this question that how much we can trust on the outcome of interpretations. For example, [46] discussed that it is important to examine the utility and robustness of the explanation in the context of medical imaging data. They posit that the explanation trustworthiness require repeatability and reproducibility. In addition, in the context of MS feature importance discovery we also posit that the explanations need to be consistent from one sample to other samples of the same group in order to guarantee the robustness of the results. We assess this assumptions by comparing the IOUs when the network is run in cross-validation mode. This experiment specifically run on the synthetic data in order to avoid problems of disentangling errors made by the model from errors made by the explanation. First, 10% of the data is left out for testing, and the rest is used for training and validation sets in five-fold cross-validation split. On every run of cross validation, network is trained on the training set, then inference is run for testing and validation set, and finally LRP interpretation is run on the predictions. The interpretations of the test set, which are generated five times over five-fold cross-validation reaches almost 99% IOU. The high level of overlapped regions demonstrate the reproducibility and repeatability of the interpretations. Likewise, the interpretations of the five validation sets over training using five-fold cross-validation reaches almost 98% IOU. This results shows the robustness of the interpretations with respect to changing the samples in the data.

These results not only justify the stability of the interpretations and the designed classification but also imply the robustness of feature selection performance.

## 4 Biomarker Detection results on Real Dataset

In this section, the performance of the proposed method is assessed on a published benchmark LC-MS dataset [4] which we refer to as real dataset. Many other Mass spectrometry datasets are available at repositories such as PRIDE or CompMS. However, the focus of this paper is to assess the feature selection on a raw LC-MS map of samples from two conditions (healthy and control) with known biomarkers presented by their *m*/*z* and RT, which is perfectly met in the selected dataset. All the parameters and hyperparameters of the model including the classification, interpretation, and feature selection parts are maintained as they were tuned on the synthetic dataset.

### 4.1 Real-Data Description

The real LC-MS dataset, consists of two groups. The first group was derived from five serum samples of healthy individuals that have been spiked with a known concentration of spike-in peptides. The second group was obtained from the serum samples only. We refer to the first and second groups as diseased and healthy, respectively. The added peptides to the diseased group are the selection of nine peptides with different concentrations to be representative of real datasets. They have predictable retention behavior and elution order that let the ground truth available in *m*/*z* and RT [4]. LC-MS acquisition yields 13 peaks from nine peptides due to the different charges. The specifications of these peaks are presented in Table 2. The concentration of 1 *pmol*/ *μL* was selected for spike-in peptides. It is common to deplete serum of high abundant proteins such that low abundant proteins can be detected. Hence, in preparation of this data, 60 *μL* of human serum of Immunoglobulin G (IgG) and Albumin was depleted. While different concentrations of spike-in peptides (0.05, 0.1, and 0.5 *pmol*/*μL*) were evaluated, a concentration of 1 *pmol*/*μL* showed the minimal intensity that would not swamp the MS signals of serum peptides in LC-MS acquisition [4] (Please see supplementary material for visualization of the spike-in peaks). We quantize the raw data and form chromatograms matrices, which are then converted into images whose width and height are *m*/*z* and RT, respectively. Each RT bin on the y-axis presents seven seconds of the MS level-1 scan, and x-axis covers ions of *m*/*z* 350 to *m*/*z* 2000. Pixel intensities are demonstrating the ion-counts. LC was run for 240 minutes, however, similar to the benchmark methods, we filter the samples to retain features within 150 minutes because there is no significant peak out of this range. We remove the features with the ion-count intensities less than two as the only noise reduction on the samples.

**TABLE 2.** Specification of the real data spike-in peptides. Base peak chromatograms of the group with spike-in peptides are presented based on their mass-to-charge ration (*m*/*z*), retention time (RT), and ion charge.

| Features No. | 1 | 2 | 3a | 3b | 4 | 5a | 5b | 6 | 7a | 7b | 8 | 9a | 9b |
| --- | --- | --- | --- | --- | --- | --- | --- | --- | --- | --- | --- | --- | --- |
| $m/z$ | 501.25 | 450.23 | 530.78 | 354.19 | 523.77 | 648.84 | 432.89 | 586.98 | 624.99 | 630.35 | 943.43 | 712.43 | 570.15 |
| Charge | 2 | 2 | 2 | 3 | 2 | 2 | 3 | 3 | 3 | 3 | 3 | 4 | 5 |
| RT(min) start-end | 4-8 | 45-49 | 53-56 | 53-56 | 59-62 | 63-67 | 63-67 | 73-77 | 77-81 | 82-86 | 79-83 | 103- 107 | 103-107 |

### 4.2 Results

Our proposed method is intended to detect differentially abundant spike-in peaks as biomarkers and to keep detected FP peaks low. We aim to decrease the amount of the FP candidates that saves lots of time for data analysis and for further examinations, but at the same time, we deem to avoid losing true positive peaks which make an important role for example in early diagnosis of diseases The evaluation will be reported as the exact number of TP and FP peaks. Table 3 compares our proposed method on the described real dataset with the benchmark methods including: msInspect, MZmine 2, Progenesis, and XCMS [4]. The first row in Table 3 demonstrates that our method outperforms the other methods in terms of detecting fewer FP peaks without being depended to the preprocessing steps used in other workflows. We follow the same statistical analysis on the selected peaks, similar to [4]. The t-test for *p* < 0.05 is calculated on each selected feature, and multiple testing correction (Benjamini-Hochberg method [47]) is applied. The features that satisfy *q* < 0.05 are selected as the discriminating features presented on the third row of Table 3. The fourth row shows the number of selected features satisfied *q* < 0.05 and fold change (FC) > 10. We detect nine biomarker peaks similar to msInspect, while we achieve almost 10 times fewer FP peaks, 195 in comparison with 2099 FP peaks in msInspect. We also outperform MZmine 2 and Prognesis with respect to both evaluation metrics, namely the number of biomarker peaks (seven in MZmine 2 and eight in Prognesis) and FP peaks (539 in MZmine 2 and 467 in Prognesis). Our experiments show that although XCMS finds fewer FP peaks, it looses low intensity 9a and 9b peaks, while DLearnMS is able to find them. Note that, as already emphasized FPs reduction should not result in loosing TPs. Specially in medical domain application, it is crucial to avoid loosing the peaks that are deemed as potential candidate for disease biomarkers. The last two columns of the Table 3 demonstrate incorporating healthy samples interpretation, 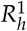, along with the diseased interpretation, 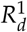. The performances show that the number of FP peaks is degraded, although it is not as pronounced as the performance on the synthetic data.

**TABLE 3.** Feature selection comparison of the proposed method with MZmine 2 [6], Progenesis LC-MS [7], and XCMS [13], which all presented in [4]. The total number of selected features is represented for all methods in the first row. Only features presented in at least two replicates in each group were used for statistical analysis for the baseline methods. The third and forth rows are demonstrating the number of features satisfying two representative criteria including t-test with multiple hypothesis testing (*q*-value< 0.05), and fold change (FC > 10). The plus sign denotes the combination of different criteria. The numbers written in parentheses indicate the selected biomarker peaks. The effect of incorporating the interpretation of diseased samples 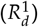 and the interpretation of healthy samples 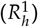 on peak detection are shown in the two last columns.

| | msInspect | MZmine 2 | Progenesis | XCMS | DLearnMS: $R_d^1$ | DLearnMS: $R_d^1, R_h^1$ |
| --- | --- | --- | --- | --- | --- | --- |
| # All selected features | 31168 (12) | 12271 (12) | 9267 (9) | 21486 (13) | 8044 (12) | 6992(11) |
| # Features for statistical analysis | 6525 (9) | 12092 (9) | 8415 (9) | 8703 (10) | 8044 (12) | 6992(11) |
| t-test ($q < 0.05$) | 4824 (9) | 3505 (7) | 4465 (9) | 1896 (7) | 3985 (11) | 3499(11) |
| t-test ($q < 0.05$) + FC ($> 10$) | 2099 (9) | 539 (7) | 467 (8) | 66 (7) | 222 (9) | 195(9) |

The biomarker peaks that are selected according to the statistical analysis are presented in Table 4. Six peaks that are commonly selected by all four other methods as differentially abundant [4] peaks have also been detected by our method.

**TABLE 4.** Real data biomarker detection comparison according to the statistical analysis. Detected differential abundant spike-in peaks are shown by check marks. Note that, our method detects all the features that are commonly selected by all other methods.

| Features No. | 1 | 2 | 3a | 3b | 4 | 5a | 5b | 6 | 7a | 7b | 8 | 9a | 9b |
| --- | --- | --- | --- | --- | --- | --- | --- | --- | --- | --- | --- | --- | --- |
| msInspect | ✓ | ✓ | ✓ | ✓ | ✓ | ✓ | ✓ | ✓ | - | ✓ | - | - | - |
| MZmine 2 | ✓ | ✓ | ✓ | ✓ | ✓ | ✓ | ✓ | - | - | - | - | - | - |
| Progenesis | ✓ | ✓ | ✓ | ✓ | ✓ | - | ✓ | - | - | ✓ | - | - | ✓ |
| XCMS | ✓ | ✓ | ✓ | ✓ | ✓ | - | ✓ | - | - | ✓ | - | - | - |
| DLearnMS | ✓ | ✓ | ✓ | ✓ | ✓ | - | ✓ | - | - | ✓ | - | ✓ | ✓ |

## 5 Conventional Machine Learning Models for high-throughput LC-MS Data Classification

In this section, we discuss the challenges that hinder classical ML methods for LC-MS data classification and why we rather use DL models in the first place. In this study, all the experiments are carried out on raw data without any dimension reduction to avoid loosing information. This might, however, cause the model to overfit or at worst cause the model performs well on training and testing but not after deployment. This is where the model’s interpretation makes roles, but not always all the ML models can be interpreted. To examplify this shortcoming, we compare the classification comparison of ML methods including, support vector machine (SVM) with linear kernel, decision tree (DT), and Adaboost with our CNN model in Table 5. The parameters of the selected methods are tuned using grid search in scikit-learn. We use five-fold and leave-on-out cross-validation for training on the synthetic and real datasets, respectively. As it is apparent from Table 5 in contrary to our designed CNN which perform equally well across two datasets, there are a huge gap in the classification performances of ML methods between the synthetic data and the real data. To investigate we tried to interpret the results and check if model make decision based on relevant discriminating features. There are model agnostic methods that enable estimating the importance of features for decision making by predictive models, such as permutation feature importance, that measures importance of features by randomly shuffling them and tracking the drop in the model’s score, or LIME [48], which locally interprets any model around a single prediction through perturbing instances and fit a linear model on the perturbations. These methods, however, are computationally infeasible for measuring the importance of high-dimensional LC-MS instances that could have more than 50000 features On the other hand, employing inherently interpretable models that enable reliable explanations are not capable of correctly classifying complex LC-MS data. For example, linear models in which the weights of the variables serve as the explanation or shallow decision trees in which the normalized total reduction of the Gini index by every feature yields the explanation. These models do not even fit on synthetic data based on our experiments. Hence, in Table 5, despite Adaboost that is not inherently interpretable and Decision tree (DT) that is not shallow enough to be interpreted, linear SVM can still be explained by the weights assigned to the features. According to this table, SVM reaches comparable classification performance as the CNN. However, the explanation results in a very poor IOU - less than 10% - between the important features selected by coefficient of SVM model and actual differences. This effect - the high accuracy and weak explanation-resulted by SVM can be explained by low fidelity of the model’s interpretation or overfitting of the model caused by some biases or pattern (comes with the simulation), unrelated to actual differences. But, the overfitting effect is more likely since SVM with the same parameter setting, trained on the synthetic data, results in a very poor accuracy on the real data. The overfitting effect can also be explained by the Adaboost and DT classification gap between the real and synthetic data as well.

**TABLE 5.** Classification comparison of the convolutional neural network (CNN) with conventional machine learning methods including, decision tree (DT), support vector machine (SVM), and adaboost. CNN shows significantly better classification performance on the real datasets. The interpretation is not available for weak classifiers. On the synthetic dataset ML methods are as accurate as CNN. However, SVM interpretation demonstrates the overfitting effect. Interpretation on the synthetic data is reported by intersection over union (IOU) between the selected and true peaks. Interpretation on the real data is reported by the amount of true positive peaks from 13 spike-in peaks. ’-’ shows no interpretation is available for the models.

| Synthetic dataset | Accuracy | Sensitivity | Specificity | Interpretation (IOU) |
| --- | --- | --- | --- | --- |
| SVM | 0.98 | 0.99 | 0.98 | feature importance(< 0.1) |
| DT | 1.0 | 1.0 | 1.0 | - |
| Adaboost | 0.99 | 1.0 | 0.99 | - |
| CNN | 1.0 | 1.0 | 1.0 | LRP (0.85) |
| Real dataset | Accuracy | Sensitivity | Specificity | Interpretation (TP/13) |
| SVM | <0.5 | <0.5 | <0.5 | - |
| DT | <0.5 | <0.5 | <0.5 | - |
| Adaboost | <0.5 | <0.5 | <0.5 | - |
| CNN | 0.8 | 0.8 | 0.8 | 12/13 |

Unreliability and poor performance of ML models on raw high-throughput LC-MS proteomics demonstrate the reasons that we choose DNN models for our analysis. We exemplify not only DNNs enable reaching the high performance, but also their interpretation is now more alleviated by the recent interpretation technologies.

## 6 Implementation Setup

The experiments in this study are implemented in Python for data analysis, Scikit-learn library [49] for ML analyzes, Keras [50] with Tensorflow backend [51] for DL analysis, and “iNNvestigate” library [52] for DL interpretation analysis on a machine with a 3.50 GHz Intel Xeon(R) E5-1650 v3 CPU and a GTX 1080 graphics card with 8 GiB GPU memory. The classification network is trained for 20 epochs and batch size of two using Adam optimizer [53] with the learning rate of 0.00001, and momentum of 0.9. We use binary crossentropy as the loss function. The kernel size in all layers is set to 3×3 with the dropout rate of 0.3. The convolution layers in the network are two dimensions and contain the following number of kernels: 32 in the first and second layers, 64 in the third layer, and two in the fourth layer. The fully connected layer as the last layer has two neurons for binary classification^1^.

## 7 Discussion

Identifying a set of biomarkers (proteins in this study) from LC-MS data is a standard task in the context of precision medicine. Performing this task on raw data is challenging due to the high dimensionality, complexity, and high noise level. Despite available tools, current workflows require several preprocessing steps to address LC-MS biomarker detection. Moreover, learning biomarkers directly using ML/DL models using supervised models require peak level annotations which can be too expensive or even infeasible to acquire. On top of it all, despite the importance of interpretable explanation in biomedical settings the application of ML/DL interpretation has been neglected in this area. To address aforementioned challenges, we introduce a deep learning (DL)-based method combined with an approach for interpretation of the learned DL configuration using the layerwise relevance propagation (LRP) technique [31]. We showed how to use the interpretation to identify potential biomarkers. Our method only requires class labels for the given training data – rather than expensive peak annotations – and is independent of otherwise necessary preprocessing steps. We trained a CNN network on the LC-MS map of the healthy and diseased samples and then used LRP interpretation firstly for network architecture selection and secondly for learning from the trained network where to look for differentially abundant peaks as biomarkers.

The first challenge with any supervised DL method is that it requires a large labeled dataset for parameter tuning; otherwise, it overfits quickly, particularly on the high dimensional and sparse LC-MS dataset. Due to the insufficient real labeled LC-MS dataset for training, our model was tuned and optimized on a large synthetically generated dataset. Besides, we verified the model robustness by measuring the dependency of the network’s decision on true features. The second challenge is that the interpretation of a DL model is not always informative when it comes to very small discriminating peaks in the sparse LC-MS dataset. Therefore, we run systematic experiments using feature selection metrics to quantitatively measure the network’s interpretation.

According to the results in Section 2.2, we showed the interpretations of different network architectures that share similar classification performance - with almost 99% training and testing accuracy - differ considerably. These differences consequently affect biomarker detection. To select network architecture we quantitatively assessed interpretation of these networks, and select the one whose interpretation is aligned the most with discriminating regions of the data. We examined the repeatability, reproducibility, and robustness of the selected model interpretation through cross validation on synthetic data in Section 3.5. Then, we built the biomarker detection on the interpretation of the selected network.

We assessed the biomarker detection of the proposed tuned model on a real dataset with predictable spike-in peptides. We showed DLearnMS achieved overall better performance in comparison with the conventional methods ([4], [6], [7], [13]) in terms of detecting fewer FP peaks despite being independent to otherwise necessary preprocessing steps.

Training the DL model on small datasets is not often recommended due to underfitting, overfitting effects, and lack of sufficient evidence (labeled data) to show the model robustness. We showed that a properly designed network can still be reliable through its validation using a proper DL interpretation.

On the synthetic data, we showed that exploiting the interpretation of *both classes* can considerably improve the FP in comparison with the setting when only the diseased class were considered. This observation stressed the importance of understanding the implications that are provided by interpretation analyzes. Leveraging this valuable information can foster more plausible network architectures resulting in a more meaningful conclusion. Recent advances in the image processing field confirm this important fact [26], [31], [54].

The improvement in the FP rate on the real dataset was not as pronounced as the synthetic dataset. This behavior can be statistically explained by the number of samples in the synthetic dataset (~ 8000) that outnumber the real dataset (~ 10). We calculated the interpretation analysis on the mean of the samples’ intensities. Therefore, the mean intensities on the large set of data is a better representative of whole data distribution than a small set. Consequently, the importance of features belonging to the larger dataset, which are assigned by the network’s decision, would be more precise.

According to Section 5, conventional ML models are failed to correctly fit on LC-MS real dataset. Despite high accuracy on the synthetic data, the poor interpretation of linear SVM on synthetic data and the huge gap between classification performance of real and synthetic data demonstrate the overfitting effect.

This study was assessed on the dataset whose biomarkers have been spiked before LC-MS acquisition. To further our research, we plan to apply our proposed method to real diseased cases. This study can be extended to the multi-subject localization of biomarkers. In this case, the interpretation of a robust multi-class classification network on the LC-MS map of samples would highlight the dominant differences of each class from the others. These differences are the potential position of biomarkers. We also consider adapting different LRP rules to different layers of the network due to their confirmed success in machine vision applications [26].

## 8 Conclusion

We present DLearnMS, an interpretable deep learning approach for LC-MS biomarker detection. DLearnMS is built on a generalized convolutional neural network combined with an interpretation method to allow understanding of the results. We successfully leverage the quantification of deep learning prediction interpretations for biomarker identification. Towards this end, the lack of labeled LC-MS data is addressed by utilizing synthetically generated data for model parameter tuning and optimization of the network architecture. DLearnMS shows bette results compared to conventional biomarker detection methods (such as msInspect, MZmine 2, Progenesis, and XCMS) in terms of detecting fewer false positive peaks and maintaining true positives – while decreasing additional computational costs by excluding commonly used preprocessing steps.

## Supporting information

This file contains supplementary material of our paper, mostly dedicated to the further simulation analyses.

## Acknowledgment

This work was supported by the German Ministry for Education and Research (BMBF) as Berlin Big Data Center (01IS14013A) and the Berlin Center for Machine Learning (01IS18037I) and within the Forschungscampus MODAL (project grant 3FO18501).

---

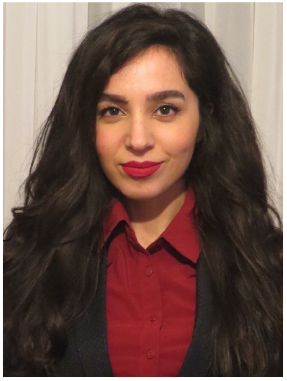

**Sahar Iravani** received her B.Sc. and M.Sc. degrees both in electrical engineering from Tabriz University, and Babol Noshirvani University of Technology, Iran, in 2012 and 2015, respectively. She is currently pursuing her PhD studies in bioinformatics at Freie Universität Berlin, Germany. She is also a working as a research assistant at Zuse institute Berlin. Her research interests lie at the intersection of machine learning, deep learning, interpretability, and high-throughput biomedical data analysis with a specific interest in mass-spectrometry proteomics analysis.

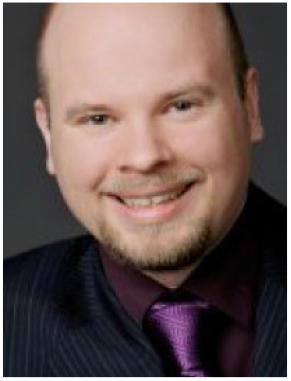

**Tim O.F. Conrad** is head of the Bioinformatics in Medicine group at Zuse Institute Berlin (ZIB) and principal investigator at the Berlin Institute for the Foundations of Learning and Data (BIFOLD). He studied Bioinformatics and Computer Science in Berlin and Melbourne. His research interests include machine learning for bio-medical data and network-based data integration.

## Footnotes

1 The datasets and implementation are available upon request from the first author.

