## Supplementary material for "An Interpretable Deep Learning Approach for Biomarker Detection in LC-MS Proteomics Data": This file contains supplementary material of our paper, mostly dedicated to the further simulation analyses.

In this Supplement, we provide the information of the peptides in synthetic and real datasets in Appendix A. In Appendix B The classification network convergence on the synthetic and real datasets are visualized. In Appendix C the distributions of real data instances around spik-in peptides are depicted.

### Appendix A
### Peptides in Synthetic dataset

TABLE 1
The accession number associated with diseased and healthy group in the synthetic dataset.

| Classes | Peptide sequences |
|---|---|
| Healthy | Q9NYW0, Q9NYV9, P59538, P59539, Q96CE8, Q96A56, O75478, Q86TJ2, Q15543, Q15573, Q9H5J8, O00268, Q9UI15, Q9H2K8, Q17R31, P10636, P68366, A6NHL2, Q13509, Q9NVG8 |
| Diseased | Q9NYW0, Q9NYV9, P59538, P59539, Q96CE8, Q96A56, O75478, Q86TJ2, Q15543, Q15573, Q9H5J8, O00268, Q9UI15, Q9H2K8, Q17R31, P10636, P68366, A6NHL2, Q13509, Q9NVG8, Q9HA65, Q9ULP9 |

TABLE 2
Real data spike-in peptide sequences

| Peptide No. | Peptide sequences | charge |
|---|---|---|
| 1 | RGDSPASSKP | 2 |
| 2 | DRVYIHP | 2 |
| 3a | RPPGFSPFR | 2 |
| 3b | RPPGFSPFR | 3 |
| 4 | DRVYIHPF | 2 |
| 5a | DRVYIHPFHL | 2 |
| 5b | DRVYIHPFHL | 3 |
| 6 | DRVYIHPFHLLVYS | 3 |
| 7a | WLTGPQLADLYHSLMK | 2 |
| 7b | WLTGPQLADLYHSLMK | 3 |
| 8 | YPIVSIEDPFAEDDWEAWSHFFK | 3 |
| 9a | GIGAVLKVLTTGLPALISWIKRKRQQ | 4 |
| 9b | GIGAVLKVLTTGLPALISWIKRKRQQ | 5 |

### Appendix B
### Visualize Convergence Distribution

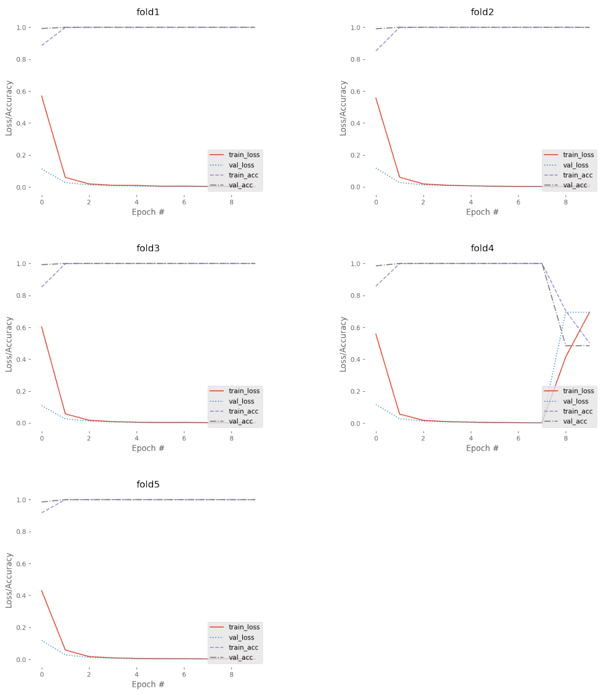


Fig. 1. Training on the simulated data. Training and validation classification accuracy are shown in purple dash line and back dash-doted line, respectively. Training and validation losses are also shown along with the accuracies in red line and blue dotted line. In this plot, we demonstrate the five fold cross-validation training curves for 10 epochs. However, for the classification comparison and continue with interpretation and feature selection the early stopping has been considered. Therefore, training is stopped after five epochs which avoided the divergence on the forth fold.

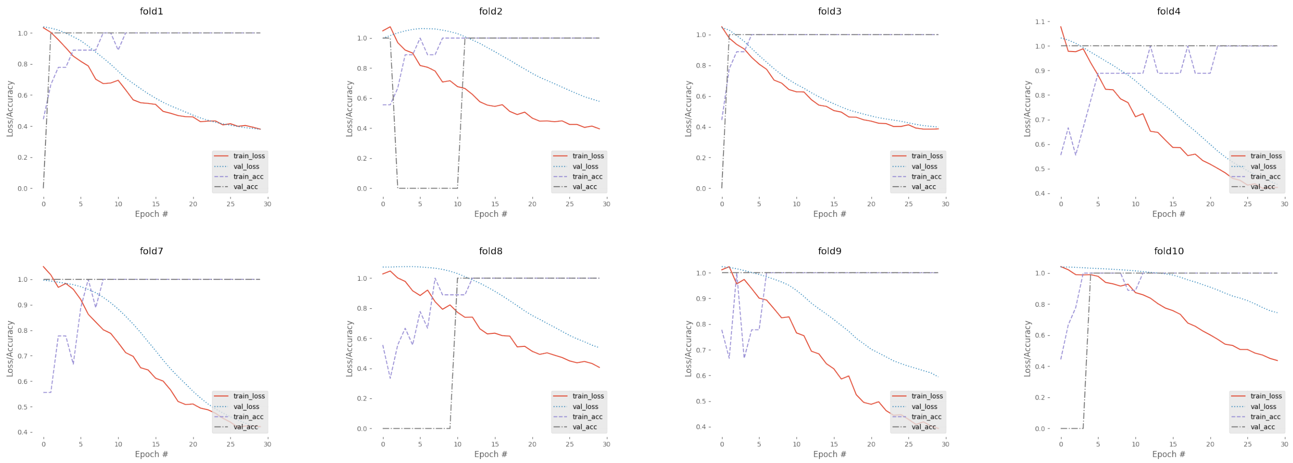


Fig. 2. Training on the real data. Training and validation classification accuracies are shown in dash line and back dash-doted line, respectively. Training and validation losses are also shown along with the accuracies in red line and blue dotted line. The trends are less smooth than simulated data, because of smaller amount of data points in the real dataset than simulated dataset.

### Appendix C
### LC-MS Real data spike-in peptide visualization

Fig. 3. LC-MS Real data spike-in peptide visualization. Chromatograms are zoomed into the location of spiked-in peptides in the group of serum samples mixed with the spiked-in peptides (MP) and group of serum samples only (noMP)

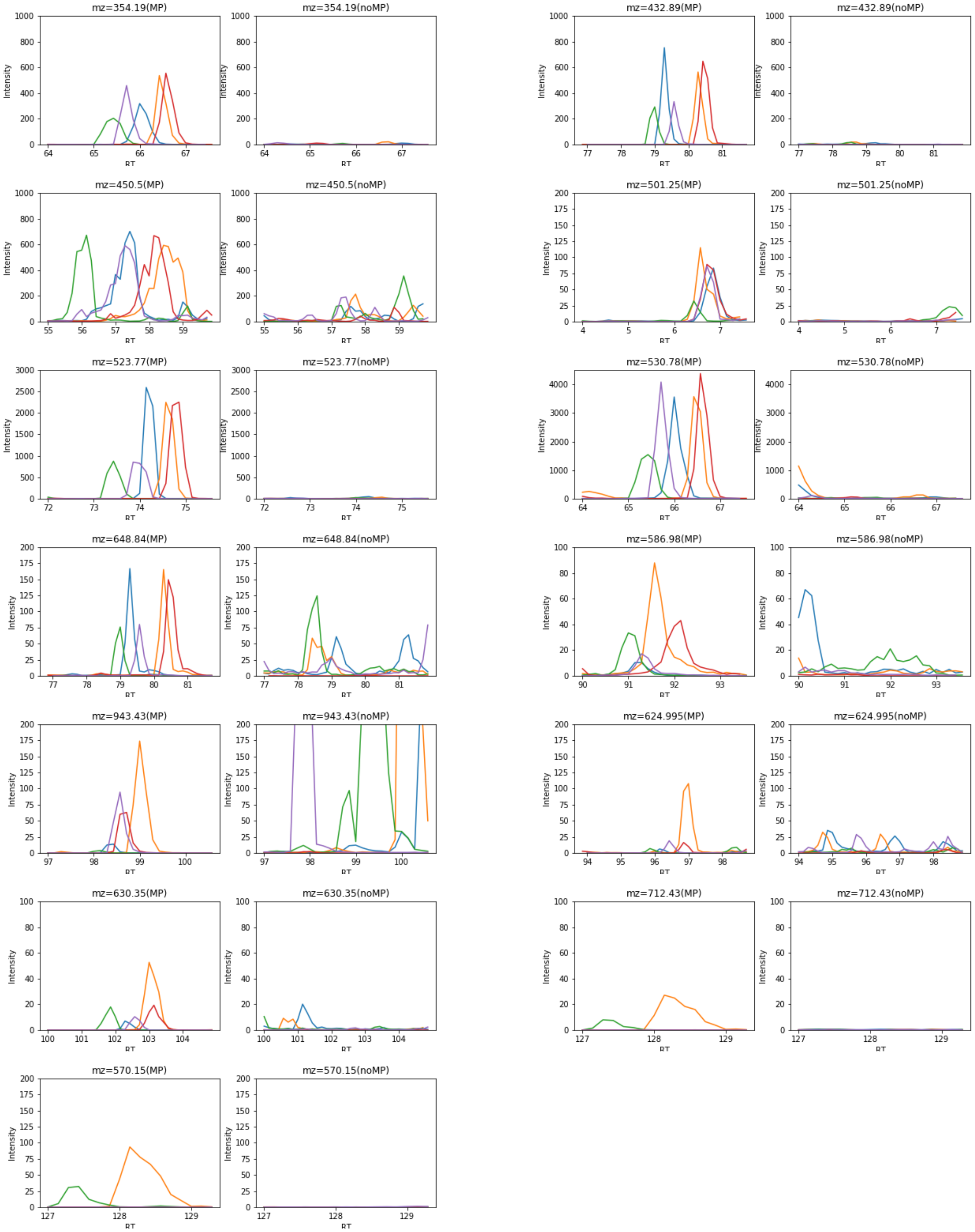
